# Asymmetric population structure in *Neoparamoeba perurans* and its kinetoplastid symbiont

**DOI:** 10.64898/2026.05.26.727903

**Authors:** Brendan A. Robertson, Bachar Cheaib, Konstantin Chekanov, Tina Oldham, Kimberley McKinnell, Michael Hill, Yee Wan Liu, Martin S. Llewellyn, Kyrie Dickson

## Abstract

Secondary endosymbioses typically involve photosynthetic gain-of-function, at least initially. For *Neoparamoeba perurans*, the agent of Amoebic Gill Disease in Atlantic salmon, the evolutionary significance of its non-photosynthetic kinetoplastid endosymbiont, *Perkinsela*, remains a mystery. While such endosymbionts usually mirror organellar uniparental inheritance, departures from strict vertical transmission can significantly impact holobiont evolution and the spread of traits like drug resistance. However, genomic analysis of *N. perurans* is hampered by continual traffic of bacteria in and out of the amoeba, limiting our understanding of its population dynamics. Here, we developed a dual-target AmpSeq panel from a draft *N. perurans* genome, enabling simultaneous genotyping of host and symbiont directly from xenic biological samples. Analysis of 58 North Atlantic isolates revealed a striking asymmetry in population structure: the amoeba host forms a diverse, panmictic population with low linkage disequilibrium, whereas *Perkinsela* exhibits lower diversity and strong clonality. While cophylogenetic analyses confirm a global signal of vertical fidelity, fine-scale genotyping reveals frequent shuffling of host-symbiont pairs. These results suggest that strict host-symbiont co-inheritance is not absolute, perhaps reflecting a system characterised by long-term vertical stability punctuated by intermittent reassortment. Ultimately, we validate AmpSeq as a robust, scalable tool for decoupling complex host-symbiont trajectories and monitoring multi-partner disease dynamics in aquaculture environments.

## Introduction

Symbiotic interactions are fundamental drivers of microbial evolution and ecological adaptation. In tightly integrated associations, symbionts are often inherited alongside host lineages, resulting in strong phylogenetic congruence and long-term co-diversification at the species level **(McFall-Ngai *et al*., 2013; Bright & Bulgheresi, 2010)**. However, even in predominantly clonal systems, this coupling can be eroded by introgression and lineage reassortment, generating discordant evolutionary signals despite obligate dependency **(Drew *et al*., 2021)**. Such ‘leaky’ co-inheritance challenges the assumptions of strict vertical transmission of symbiont lineages with potentially important evolutionary and adaptive consequences. For example, leakage may allow the host lineage to escape Muller’s ratchet ‘by proxy’ and purge deleterious mutations from associated symbionts, or even to benefit from potentially advantageous mutants that occur within other symbiont lineages (e.g., where a drug resistance allele might be symbiont encoded **(Tan, *et al*., 2021)**.

The marine amoeba *Neoparamoeba perurans*, the causative agent of Amoebic Gill Disease (AGD) in Atlantic salmon (*Salmo salar L*.) aquaculture **(Young *et al*., 2007)**, harbours an obligate intracellular kinetoplastid endosymbiont of the genus *Perkinsela*. Genomic analyses have demonstrated extensive metabolic integration between the host and symbiont in sister species *N. permaquidensis* **(Tanifuji, *et al*., 2017)**. Phylogenetic analyses across multiple *Neoparamoeba* species have revealed strict congruence between host and symbiont phylogenies, suggesting ancient co-evolution and vertical transmission **(Young, *et al*., 2014; Tanifuji *et al*., 2011)**. However, the evolutionary relationship between the *Neoparamoeba* host and *Perkinsela* symbiont populations within species remains unknown, particularly regarding their reproductive mode and whether *Perkinsela* is strictly vertically inherited or subject to horizontal exchange between host lineages.

*N. perurans* represents a challenge to genome biologists. Its cytoplasm is packed with commensal microbes, and the amoeba cannot be grown in axenic culture **(MacPhail, *et al*., 2021)**. To date, there is no available reference genome for this important aquaculture pathogen, and population genetic studies are limited to a handful of genetic markers (ribosomal or targeted loci) **(Hansen, *et al*., 2019; Young, *et al*., 2008; Young., *et al*, 2007)**. To recover *N. perurans* SNP variation from biological samples and resolve *N. perurans* population genetic structure, we first generated a heavily contaminated draft genome sequence from xenic culture and used resultant contigs to develop a novel AmpSeq marker panel for simultaneous population genomic analysis of *N. perurans* and *Perkinsela* symbiont diversity. We genotyped isolates from Scottish and Norwegian aquaculture sites to: **(1)** quantify reproductive strategies and clonal diversity of both partners, **(2)** assess population structure and gene flow patterns across geographical scales, and **(3)** test for phylogenetic congruence indicative of strict vertical inheritance versus horizontal symbiont exchange. Our analyses reveal unexpected asymmetries in population structure and frequent disruption of clonal co-inheritance, providing insights into the evolutionary dynamics of obligate endosymbiosis and challenging traditional models of symbiont transmission fidelity.

## Materials and Methods

Isolates of *Neoparamoeba perurans* and their associated *Perkinsela* endosymbiont were obtained from AGD-affected Atlantic salmon aquaculture sites in Scotland and Norway **(Fig 1A)**. An AmpSeq panel of 55 markers was designed based on a draft of the *N. perurans* genome generated using a combination of short and long-read sequencing **(SD1)**. Both nuclear and mitochondrial loci of both the amoeba and symbiont were targeted. Following amplification, amplicon pools were sequenced via the 2x150bp Illumina platform. Reads were mapped to a *N*.

**Figure 1:**
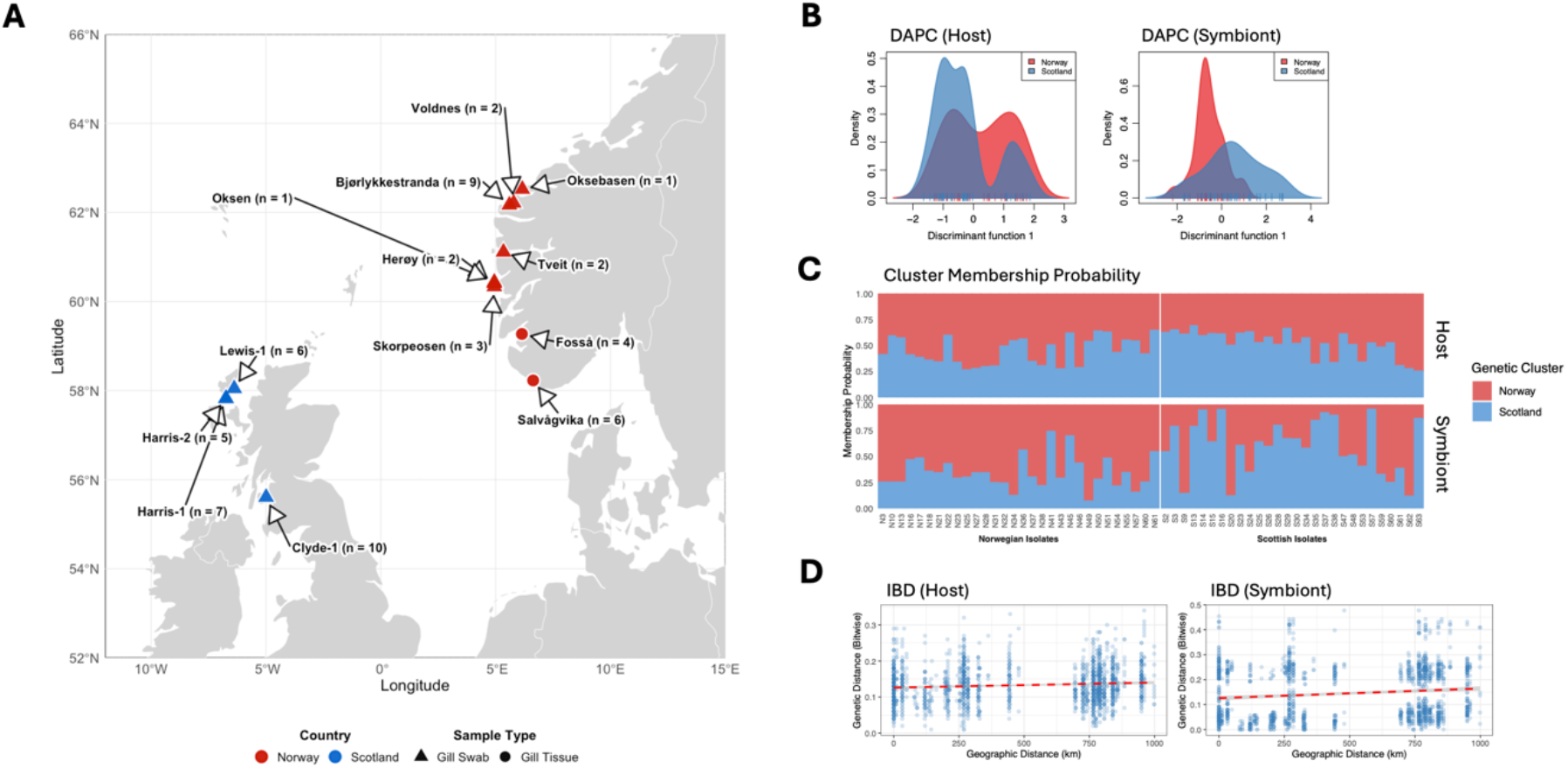
Genetic structure of host and symbiont across the North Atlantic and North Sea. (A) Geographical distribution of sampling sites. The map indicates locations of 13 Atlantic salmon aquaculture sites sampled for *N. perurans* and *Perkinsela* genomics across Norway (red symbols) and Scotland (blue symbols), spanning approximately 996 km across the North Sea and North Atlantic regions. Shape indicates sample type obtained at each location (triangles = gill swabs; circles = gill tissue). Sample sizes per site are indicated in parentheses (n = x). **(B) Discriminant Analysis of Principal Components (DAPC)** scatterplots for *Neoparamoeba* (50 SNPs), showing extensive overlap between Scotland (blue) and Norway (red) isolates, and *Perkinsela* (44 SNPs), showing more discrete clustering by region. To prevent model overfitting, the optimal number of retained principal components was determined via a-score optimisation (1 PC for host, 5 PCs for symbiont). **(C) Cluster Membership Probabilities** (K=2) for host and symbiont. High levels of admixture in the host support panmixia *(F*_*ST*_ *= 0*.*008, p = 0*.*123)*, while the symbiont shows a higher and significant degree of lineage assignment to geographic clusters *(F*_*ST*_ *= 0*.*035, p < 0*.*001)*. **(D) Isolation by Distance (IBD)** plots showing the relationship between geographic distance (km) and bitwise genetic distance. Whilst the regression slopes for both the *Neoparamoeba* host and *Perkinsela* symbiont are significant, the low r values indicate that geographic distance explains ∼1% of the observed genetic variation *(r*_*host*_ *= 0*.*09, r*_*symbiont*_ *= 0*.*12)*.

*perurans* draft reference genomes using BWA-MEM, and SNPs were jointly called using bcftools under a multiallelic model. Low-confidence genotypes were masked, and variants were filtered based on quality and missingness thresholds for high-confidence SNP datasets **(SD1)**.

Population genomic analyses were conducted **(SD1)**. Population structure was assessed using discriminant analysis of principal components (DAPC), reproductive mode was evaluated using multi-locus genotype clustering and Index of Association, and genetic distance between isolates was visualised using minimum spanning networks (MSN). Isolation by distance (IBD) was assessed using Mantel tests, while host-endosymbiont evolutionary coupling was assessed using both Mantel tests and a Procrustes approach of co-phylogeny (PACo), with association structure and phylogenetic incongruence further visualised using bipartite host-endosymbiont MLG networks.

## Results

### Optimisation of AmpSeq Panel for Marine Holobionts

Following stringent quality control, the AmpSeq panel successfully resolved 50 high-confidence SNPs for the *Neoparamoeba* host and 44 for the *Perkinsela* symbiont across 58 isolates, achieving deep coverage (median depth >150x) capable of supporting robust micro-evolutionary analyses **(SD1 for further details)**.

### Asymmetric Population Structure and Dispersal Dynamics

Population genomic analyses revealed asymmetry in spatial structuring between the two symbiotic partners. The *Neoparamoeba* host population exhibited a largely panmictic signature across the North Atlantic (FST = 0.008; p = 0.123) with extensive overlap in DAPC density distributions **(Fig 1B)**. Cluster membership probabilities further supported this pattern, revealing largely undigerentiated ancestry proportions across sites in the host **(Fig 1C)**. Conversely, *Perkinsela* lineages exhibited a weak but significant geographic signal (FST = 0.035; p < 0.001) **(Fig 1B)**, suggesting the symbiont is more sensitive to dispersal barriers than its host. Whilst isolation-by-distance (IBD) analysis was statistically significant for both partners (*r*_*Host*_ = 0.091, p = 0.001; *r*_*Symbiont*_ = 0.120, p = 0.001; **Fig 1D**), low correlation coefficients indicate that geographic distance explains only ∼1% of genetic variance. This suggests that extensive gene flow - likely facilitated by the movement of aquaculture stock and ocean currents - overwhelms local genetic isolation.

### Divergent Clonal Architecture and Genomic Contraction

Both partners exhibited significantly clonal reproductive modes under outbreak conditions (Index of Association, *p = 0*.*001;* **SD2 Fig S2A**), yet on different evolutionary trajectories. The host maintained higher genetic diversity, represented by 47 distinct multi-locus genotypes (MLGs) **(Fig 2A)**, and low pairwise linkage disequilibrium (*r*^*2*^ *> 0*.*2*, ∼5% **(Fig 2B)**, consistent with periodic recombination. In contrast, the symbiont appears locked in a tighter clonal trajectory, contracting into just 21 MLGs **(Fig 2A)** with a four-fold higher frequency of high-linkage SNP pairs (∼20%) (**Fig 2B**). This pattern likely reflects the genomic contraction and increased genetic drift typical of obligate endosymbionts.

**Figure 2:**
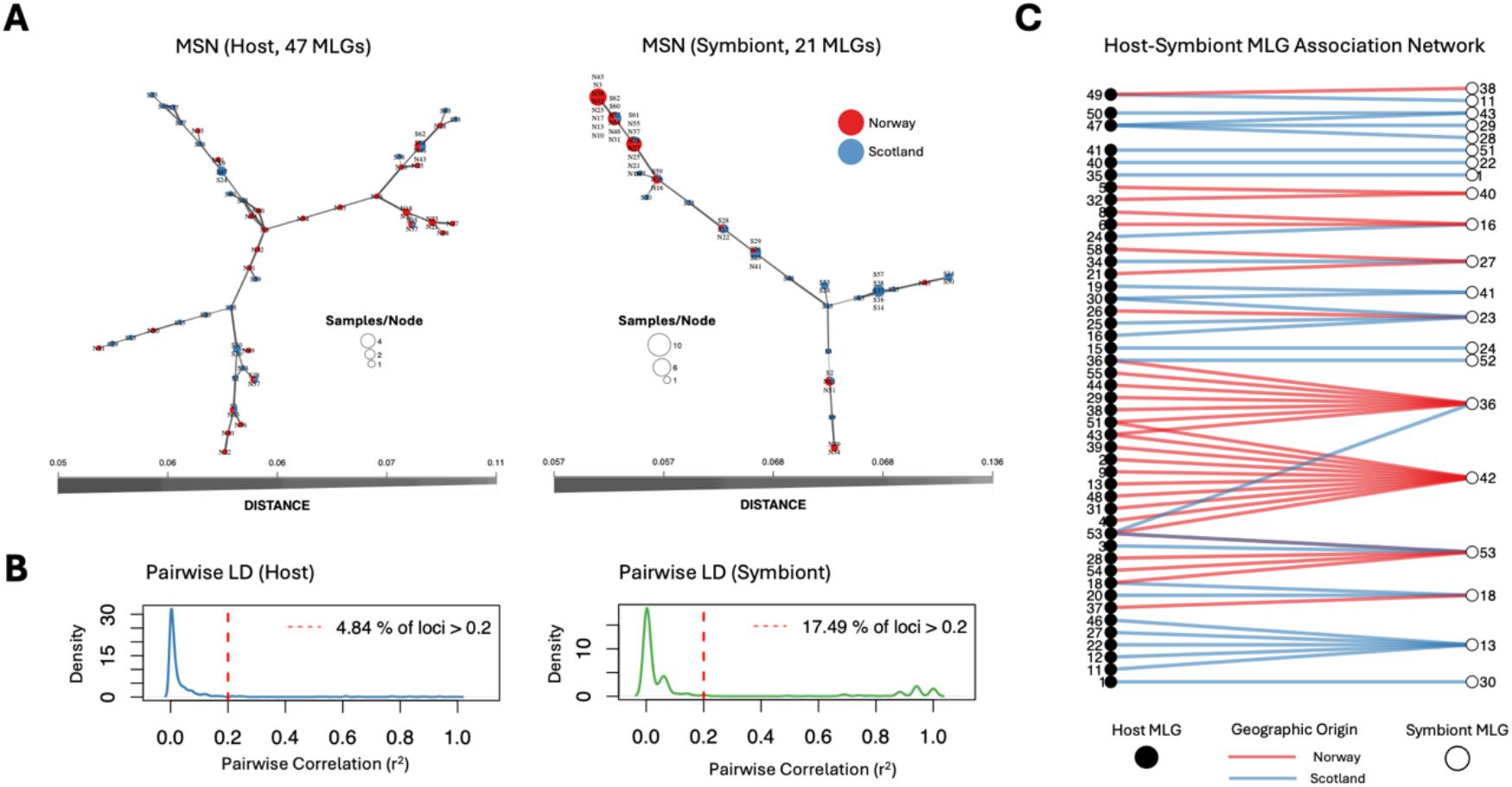
Host-Symbiont co-evolution and transmission dynamics. **(A) Minimum Spanning Network (MSN)** based on bitwise genetic distance. The highly reticulated topology of the *Neoparamoeba* host is indicative of recombination and high genotype diversity (47 Multi-Locus Genotypes, MLGs). In comparison, the *Perkinsela* symbiont MSN demonstrates a clonal topology where multiple isolates collapse into 21 distinct MLGs. MSN node size corresponds to the number of isolates sharing a genotype; edge thickness represents genetic similarity; colour-coding of MSN node represents the proportion of isolates within each MLG belonging to either Norway (red) or Scotland (blue). **(B) Genome-wide Pairwise Linkage Disequilibrium (LD)**. Density distributions of the squared Pearson correlation coefficient *(r*^*2*^) for all pairwise combinations of SNPs across the host (blue) and symbiont (green) genomes. Pairwise *r*^*2*^ values were calculated globally (ignoring physical genomic distance) to assess overall genome clonality. The red dashed vertical line indicates a widely accepted threshold for biologically meaningful LD *(r*^*2*^ *> 0*.*2)*. The host *Neoparamoeba* exhibits a relatively recombining population structure, with only 4.84% of SNP pairs demonstrating significant linkage *(r*^*2*^ *> 0*.*2)*. In contrast, the *Perkinsela* symbiont displays a highly clonal signature, with 17.49% of genome-wide SNP pairs in linkage disequilibrium. **(C) Bipartite association network of host and symbiont MLGs**. Circles represent unique MLGs for *Neoparamoeba* (black) and *Perkinsela* (white), with adjacent numbers denoting specific MLG identities. Connecting lines indicate observed pairings, coloured by geographic origin (red: Norway; blue: Scotland). Whilst the majority of the population follows vertical inheritance, several host MLGs (e.g., 47, 30, 53) were found to harbour several different symbiont MLGs.

### Cophylogenetic Congruence and Transmission Dynamics

Global association tests indicated that the host and symbiont populations are not genetically independent (Mantel *r = 0*.*130; p = 0*.*049)*. Procrustes superimposition (PACo) demonstrated a highly significant global topological association *(m*^*2*^ *= 14*.*3; p < 0*.*001;* **SD2 Fig S2B)**, supporting a primary mode of vertical transmission. This was corroborated by ParaFit *(Global Stat = 5*.*23; p = 0*.*039)*, though the marginal p-value suggests the presence of substantial localised incongruence.

Fine-scale assessment of clonal co-inheritance revealed breakdowns in strict verticality; 8 host MLGs harboured non-matching *Perkinsela* MLGs **(Fig 2C)**. This ‘leaky’ signal was further highlighted by PACo squared residuals, which identified specific isolates (e.g., S9, N51) as high-incongruence outliers **(SD2 Fig S2C)**. Conversely, single symbiont lineages were observed colonising multiple divergent host backgrounds **(Fig 2C)**. This bidirectional incongruence suggests that the *Neoparamoeba-Perkinsela* holobiont is not a closed system; rather, it follows a model of vertical transmission punctuated by host-switching, environmental acquisition, or uniparental inheritance. Such shuffling of host-symbiont pairings may be facilitated by the high-density environment of aquaculture pens, providing a mechanism for genetic reshuffling in an otherwise clonal system.

## Discussion

Genomic resources for *Neoparamoeba* remain limited, largely due to the technical difficulties of axenic culture and high bacterial contamination of environmental samples **(Young *et al*., 2007; Young *et al*., 2008)**. These challenges have historically constrained our ability to address fundamental epidemiological questions regarding *N. perurans* and its endosymbiont *Perkinsela*. Here, we demonstrate that our dual-target AmpSeq approach can successfully resolve the population structure of both the amoeba host and its endosymbiont directly from xenic North Atlantic samples. With a small SNP panel, we detected a clear biological signal: pronounced asymmetry in the evolutionary trajectories of the two partners. The host population is characterised by high genotypic richness, weak geographic structure and weak pairwise linkage disequilibrium, while *Perkinsela* exhibits lower diversity and stronger clonal structure, with a four-fold increase in linkage patterns indicative of genetic contraction and drift typical of an obligate endosymbiotic lifestyle **(Young *et al*., 2014)**.

Despite the limited marker panel size, we detected significant Isolation by Distance (IBD) in both host and symbiont. The presence of IBD in *N. perurans* suggests that North Atlantic dispersal is not entirely homogenised, potentially reflecting specific genotype-environment interactions, or faster evolutionary turnover in the amoeba resulting in the rapid accumulation of mutational differences between sites **(Hellberg, 2009; Palumbi, 2003)**. Our data suggest that *N. perurans* maintains a predominantly clonal expansion phase, likely punctuated by episodic recombination, a strategy common among facultative sexual protists **(Speijer *et al*., 2015; Weedall & Hall, 2014)**. Importantly, the fact that structure was detectable with a limited SNP panel suggests that expanded panels will have sufficient power to resolve fine-scale connectivity between individual farms and production cycles. Much larger panels of markers reveal a startling lack of IBD in other key salmonid aquaculture pathogens, for example, the salmon sea louse *Lepeophtheirus salmonis*, where high gene flow impedes fine-scale molecular epidemiological studies are impeded by the sheer rate of parasite transfer **(Jacobs, *et al*., 2018; Besnier, *et al*., 2014)**.

More striking is the decoupling of host and symbiont genotypes. While global congruence metrics support a history of vertical fidelity, fine-scale mismatches were pervasive; 10 of 11 multi-isolate host MLGs harboured non-matching *Perkinsela* MLGs, suggesting that the *Neoparamoeba-Perkinsela* association is not a fully closed system. Such discordance could mirror the ‘leaky’ organellar inheritance observed in other pathogenic protists like *Plasmodium* and *Toxoplasma* **(Collier et al., 2025)** and is consistent with the genomic interdependence characterised in the Neoparamoeba-kinetoplastid symbiosis **(Tanifuji *et al*., 2017; Tanifuji *et al*., 2011)**, though the specific mechanisms in *Neoparamoeba* require further study. It remains unclear how these organisms interact to enable recombination, or what biological factors determine which *Perkinsela* lineages are retained or replaced during these events. Crucially, the ability of symbiont lineages to move between host genetic backgrounds has significant implications for AGD emergence and adaptation, as it may facilitate the rapid spread of beneficial symbiont traits such as drug resistance **(Drew *et al*., 2021; Bright & Bulgheresi, 2010)**.

Ultimately, we demonstrate that AmpSeq fills a critical gap between low-resolution markers and cost-prohibitive whole-genome sequencing for such complex, xenic amoeba. By bypassing the need for axenic culture, this framework allows for simultaneous host-symbiont surveillance. Larger SNP panels could resolve farm-to-farm transmission pathways, providing a scalable foundation for monitoring the emergence and spread of Amoebic Gill Disease (AGD) in a changing climate.

## Supporting information

Full methods Sections.

Supplemental Figure 1, Table 1, and Figure 2. Figure one is not referenced in the text - to illustrate reads per loci per site.

## Acknowledgments

This research was supported in part by research grants from the Biotechnology and Biological Science Research Council (grant numbers BB/T016280/1, BB/Y012437/1).

## References

Besnier, F., Kent, M., Skern-Mauritzen, R., Lien, S., Hemmingsen, W., Karlsen, B.O., Aspehaug, V. and Glover, K.A. (2014) ‘Human-induced evolution caught in action: SNP-array reveals rapid amphi-Atlantic spread of pesticide resistance in the salmon ectoparasite Lepeophtheirus salmonis’, BMC Genomics, 15, p. 937. Available at: 10.1186/1471-2164-15-937.

Bright, M. and Bulgheresi, S. (2010) ‘A complex journey: transmission of microbial symbionts’, Nature Reviews Microbiology, 8(3), pp. 218–230. Available at: 10.1038/nrmicro2262.

Collier, S.L., Farrel, S.N., Goodan, C.D. and McFadden, G.I. (2025) ‘Modes and mechanisms for the inheritance of mitochondria and plastids in pathogenic protists’, PLOS Pathogens, 21(1), p. e1012835. Available at: 10.1371/journal.ppat.1012835.

Drew, G.C., Stevens, E.J. and King, K.C. (2021) ‘Microbial evolution and transitions along the parasite–mutualist continuum’, Nature Reviews Microbiology, 19(10), pp. 623–638. Available at: 10.1038/s41579-021-00550-7.

Hansen, H., Botwright, N.A., Cook, M.T., Douglas, A., Downes, J., Gallagher, M.D., Ruane, N.M. and Matejusova, I. (2019) ‘Genetic diversity among geographically distant isolates of Neoparamoeba perurans’, Diseases of Aquatic Organisms, 137, pp. 81–87. Available at: 10.3354/dao03433.

Hellberg, M.E. (2009) ‘Gene flow and isolation among populations of marine animals’, Annual Review of Ecology, Evolution, and Systematics, 40, pp. 291–310. Available at: 10.1146/annurev.ecolsys.110308.120223.

Jacobs, A., De Noia, M., Praebel, K., Kanstad-Hanssen, Ø., Paterno, M., Jackson, D., McGinnity, P., Sturm, A., Elmer, K.R. and Llewellyn, M.S. (2018) ‘Genetic fingerprinting of salmon louse (Lepeophtheirus salmonis) populations in the North-East Atlantic using a random forest classification approach’, Scientific Reports, 8, p. 91. Available at: 10.1038/s41598-018-19323-z.

McFall-Ngai, M., Hadfield, M.G., Bosch, T.C.G., Carey, H.V., Domazet-Lošo, T., Douglas, A.E., Dubilier, N., Eberl, G., Fukami, T., Gilbert, S.F., Hentschel, U., King, N., Kjelleberg, S., Knoll, A.H., Kremer, N., Mazmanian, S.K., Metcalf, J.L., Nealson, K., Pierce, N.E., Rawls, J.F., Reid, A., Ruby, E.G., Rumpho, M., Sanders, J.G., Tautz, D. and Wernegreen, J.J. (2013) ‘Animals in a bacterial world, a new imperative for the life sciences’, Proceedings of the National Academy of Sciences of the United States of America, 110(9), pp. 3229–3236. Available at: 10.1073/pnas.1218525110.

MacPhail, D.P.C., Koppenstien, R., Maciver, S.K., Paley, R., Longshaw, M. and Henriquez, F.L. (2021) ‘Vibrio species are predominantly intracellular within cultures of Neoparamoeba perurans, the causative agent of amoebic gill disease (AGD)’, Aquaculture, 532, p. 736083. Available at: 10.1016/j.aquaculture.2020.736083.

Palumbi, S.R. (2003) ‘Population genetics, demographic connectivity, and the design of marine reserves’, Ecological Applications, 13(S1), pp. 146–158. Available at: 10.1890/1051-0761(2003)013[0146:PGDCAT]2.0.CO;2

Speijer, D., Lukeš, J. and Eliáš, M. (2015) ‘Sex is a ubiquitous, ancient, and inherent attribute of eukaryotic life’, Proceedings of the National Academy of Sciences of the United States of America, 112(29), pp. 8827–8834. Available at: 10.1073/pnas.1501725112.

Tan, S., Mudeppa, D.G., White, J. III, Patrapuvich, R. and Rathod, P.K. (2021) ‘Properties of Plasmodium falciparum with a deleted apicoplast DNA gyrase’, Antimicrobial Agents and Chemotherapy, 65(9), p. e00586–21. Available at: 10.1128/AAC.00586-21

Tanifuji, G., Cenci, U., Moog, D., Dean, S., Nakayama, T., David, V., Fiala, I., Curtis, B.A., Sibbald, S.J., Onodera, N.T., Matsuo, E., Jimenez, V., Handschuh, M., Suzuki, S., Kamikawa, R., Ishida, K., Inagaki, Y., Keeling, P.J., Hashimoto, T., Lukes, J. and Archibald, J.M. (2017) ‘Genome sequencing reveals metabolic and cellular interdependence in an amoeba-kinetoplastid symbiosis’, Scientific Reports, 7(1), p. 11688. Available at: 10.1038/s41598-017-11866-x.

Tanifuji, G., Kim, E., Onodera, N.T., Gilbeault, R., Dlutek, M., Cawthorn, R.J., Fiala, I., Lukeš, J., Greenwood, S.J. and Archibald, J.M. (2011) ‘Genomic characterization of Neoparamoeba pemaquidensis (Amoebozoa) and its kinetoplastid endosymbiont’, Eukaryotic Cell, 10(8), pp. 996–1006. Available at: 10.1128/EC.05027-11.

Weedall, G.D. and Hall, N. (2015) ‘Sexual reproduction and genetic exchange in parasitic protists’, Parasitology, 142(S1), pp. S120–S127. Available at: 10.1017/S0031182014001693.

Young, N.D., Crosbie, P.B., Adams, M.B., Nowak, B.F. and Morrison, R.N. (2007) ‘Neoparamoeba perurans n. sp., an agent of amoebic gill disease of Atlantic salmon (Salmo salar)’, International Journal for Parasitology, 37(13), pp. 1469–1481. Available at: 10.1016/j.ijpara.2007.04.018.

Young, N.D., Dyková, I., Snekvik, K., Nowak, B.F. and Morrison, R.N. (2008) ‘Neoparamoeba perurans is a cosmopolitan aetiological agent of amoebic gill disease’, Diseases of Aquatic Organisms, 78(3), pp. 217–223. Available at: 10.3354/dao01869.

Young, N.D., Dyková, I., Crosbie, P.B.B., Wolf, M., Morrison, R.N., Bridle, A.R. and Nowak, B.F. (2014) ‘Support for the coevolution of Neoparamoeba and their endosymbionts, Perkinsela amoebae-like organisms’, European Journal of Protistology, 50(5), pp. 509–523. Available at: 10.1016/j.ejop.2014.07.004

