## Supplementary material for "Asymmetric population structure in *Neoparamoeba perurans* and its kinetoplastid symbiont": Full methods Sections.

### Supplementary Data 1 (SD1) - Methods Section

#### S1. Sample Collection and Sample Sites

AGD-affected samples were obtained from Patogen belonging to Atlantic salmon producers across the North Atlantic region between 2021 to 2023. Norwegian samples were obtained from nine aquaculture sites operated by Mowi (2021-2022), accessed through the Norwegian Institute of Marine Research, spanning locations from Fossá in the south to Bjerlikkestranda in the north. Scottish samples were obtained from four sites operated by Bakkafrost (2023) across the Hebridean islands Lewis and Harris, and the firth of Clyde. Biological samples consisted of gill swabs and gill tissue (only Fossá and Salvågvika) collected from clinically affected fish exhibiting signs of AGD. Site co-ordinates were extracted from metadata and visualised in R using **ggplot2 (Wickham, 2016)**, **sf**, and **rnaturalearth** packages **(Massicotte & South, 2023; Slowikowski, 2023; Pebesma, 2018)**. The sampling area spanned approximately 996km across the North Sea and North Atlantic, encompassing a latitudinal range from 55.6˚N to 62.5˚N and a longitudinal range from 6.7˚W to 6.6 ˚E. The spatial distribution of 13 sampling sites was used to capture regional population structure across major Atlantic salmon aquaculture regions.

#### S2. DNA extraction

DNA extractions were conducted on Atlantic salmon gill swabs and gill tissue obtained from Norway (n = 10-gill tissue, n = 20-gill swabs) and Scotland (n = 28-gill swabs) using a Qiagen DNeasy Blood & Tissue kit (Qiagen, Doncaster, VIC, Australia) following the manufacturer's standard protocol. All samples were quantified for DNA concentration (ng/µl) and purity results (260/280 & 260/230 ratios), with additional quantification on Qubit High-sensitive double-strand DNA for a subset of samples.

#### S3. Positive Neoparamoeba perurans samples

Original cultures of *P. perurans* were obtained from the University of the West of Scotland and isolated as described in MacPhail *et al.* (2021). New cultures were established at the University of Glasgow in malt yeast broth (MYB) and maintained at a salinity of 35 parts per thousand (ppt). Amoebae were filtered to remove biofilm build-up using a 15µm pluriStrainer® cell strainer (PluriSelect Life Science, Leipzig, Germany) following standard protocol. To limit bacterial contamination, further filtration was conducted using a 1µm filter to obtain a pure culture of *P. perurans* at 15,000 cells/ml. DNA was extracted using a Qiagen DNeasy Blood & Tissue kit (Qiagen, Doncaster, VIC, Australia) following the manufacturer's standard protocol. Polymerase chain reaction (PCR) was conducted on the filtered positive control of *P. perurans* using oligonucleotide primers specifically designed to target the 18S rRNA gene sequence from Young, *et al.,* (2008) to amplify a 636bp fragment (Table S1). Each PCR reaction (10µl) contained 1µl of *P. perurans* control and 9µl of master mix containing: 5µl of Q5, 2µl and 1µl of each forward and reverse primer (Young_f & Young_r). The reaction was conducted using a Primer Thermal cycler (Cole-Parmer, St. Neots, Cambs, UK) with a thermal profile of 94˚C for 3 minutes, followed by 35 cycles of 94˚C for 30s, 60˚C for 30s, 72˚C for 30s and a final step of 72˚C for 10 minutes to allow the completion of any partial copies. Gel electrophoresis was then run on a 1% gel at 100V for 60 minutes to determine the presence of *N. perurans*.

**Table S1: Screening PCR primer pair for detection of *Neoparamoeba perurans*.**

| Protocol | Gene | Target | Primer | Name | Sequence (5' – 3') | Product size (bp) | Reference |
|---|---|---|---|---|---|---|---|
| PCR | 18S rRNA | *N. perurans* | Forward | Young_f | ATCTTGACYGGTTCTTTCGRGA | 636 | Young et al., 2008 |
| | | | Reverse | Young_r | ATAGGTCTGCTTATCACTYATTCT | | |

#### S4. Genomic analysis for Primer Design

##### Sequence Data Processing and Assembly

Raw Illumina sequencing data from three independent runs (Run1: AmoebaBach2 250bp paired-end; Run2 and Run3: G96 250bp and 150bp paired-end, respectively) were quality-filtered using BBDuk v38.90 **(Bushnell, 2014)** with the following parameters: minimum average quality score (maq=20), entropy threshold (0.5), maximum N bases (maxns=1), and minimum read length (100bp). Quality assessment was performed using FastQC v0.11.8 **(Andrews, 2010)** before and after trimming. Paired-end reads were merged using BBMerge **(Bushnell, 2014)** with a minimum insert size of 50bp (25bp for 150bp reads) to generate consensus sequences where possible.

##### Reference-based Read Mapping and Contamination Removal

Reference genomes for six target organisms were prepared: *Neoparamoeba pemaquidensis* (GCA_002151225.1), *Veramoeba* sp. (GCA_020696425.1), *Bodo saltans* (GCA_001460835.1), *Acanthamoeba castellanii* (GCA_000313135.1), *Paratrypanosoma* sp. (GCA_002921335.1), and *Perkinsela* sp. (GCA_001235845.1) obtained from **VEuPathDB** (Amos, *et al.*, 2022). Genomic sequences were processed using any2fasta (Seemann, 2013) to standardise formatting, followed by removal of ambiguous nucleotides (N bases) using custom AWK /shell scripts. Cleaned reads were mapped to the combined reference database using minimap2 v2.24 **(Li, 2018)** with short-read alignment parameters (-ax sr) and 96 threads, with output sorted and converted to BAM format using samtools v1.15 **(Li *et al.*, 2009)**. Mapped reads were extracted using samtools bam2fq to enrich for target organism sequences while removing host and environmental contamination. Additional contamination screening was performed using DeconSeq **(Schmieder & Edwards, 2011),** where specified.

##### Genome Assembly and Hybrid Approach

Individual assemblies were generated using MEGAHIT v1.2.9 (Li et al., 2015). For the hybrid assembly approach, long-read Oxford Nanopore data were quality-filtered using NanoFilt **(De Coster *et al.*, 2018)** with minimum length (500bp), head/tail cropping (10bp each), and minimum quality score (Q10). Long reads were mapped to reference genomes using minimap2 **(Li, 2018)** with Oxford Nanopore parameters (-ax map-ont for genomic reads, -ax splice for cDNA reads). Hybrid assemblies incorporating both Illumina short reads and Nanopore long reads were generated using Flye v2.9 **(Kolmogorov *et al.*, 2019)** (--nano-hq mode) and MaSuRCA v4.0.4 **(Zimin *et al.*, 2013)**, combining all sequencing runs and mapped reads with reference contigs to improve assembly contiguity and completeness.

##### Assembly Quality Assessment

Assembly completeness was evaluated using EukCC v2.1.1 (Saary et al., 2020) with the protozoa database, specifying relevant taxonomic IDs (630701, 1027875, 1641714, 325089, 200887) and parameters optimised for single-cell eukaryotes (marker_prevalence=90%, set_size=15, max_set_size=1000). Additional quality assessment was performed using BUSCO v5.4.3 **(Manni *et al.*, 2021)** with the Euglenozoa lineage database (euglenozoa_odb10) from OrthoDB **(Kriventseva *et al.*, 2019)**, employing both MetaEukaryotic and AUGUSTUS gene prediction methods. For AUGUSTUS analysis **(Stanke *et al.*, 2008; Stanke & Morgenstern, 2005)**, a custom species model for *Perkinsela* was trained and applied to improve gene prediction accuracy for the symbiont sequences.

##### Primer Design and Target Identification

The resulting high-quality assemblies and enriched sequence datasets from this pipeline were subsequently used to identify unique genomic regions specific to *Neoparamoeba perurans* and its intracellular *Perkinsela* symbiont for the development of species-specific PCR primers, enabling sensitive detection and quantification of these organisms in environmental and host samples.

The pipeline consisted of culture-free genome-wide locus sequence typing (GLST) using a similar pipeline published by our laboratory **(Schwabl, *et al.*, 2020)**. The pipeline consisted of mapping reads to identify unique genomic regions specific to *Neoparamoeba perurans* and its intracellular *Perkinsela* symbiont. After the mapping and SNP variants identification and quality filtration, the genomic fragments containing SNP candidates from the references were extracted, and Batchprimer3 v1.0 was used to design primers and add tags for barcoding. Afterwards, MULTIPLX v2.1.4 was used to calculate primer alignment energies and remove primers with high non-target affinity.

##### S5. Individual primer testing

Initially, PCR was conducted to determine the amplification of each of the 55 experimental target amplicons. PCR assays were performed using NEB Q5 High-fidelity colourless solution of Taq DNA polymerase (New England Biolabs, Hitchin, Herts, UK). Each reaction (10µl) contained 0.5µl of *N. perurans* control and 9µl of master mix containing: 5µl of Q5, 3.5µl of Nuclease-free water and 1µl of each primer pair. The reaction was conducted using a Prime Thermal cycler (Cole-Parmer, St. Neots, Cambs, UK) with a thermal profile of; 98°C for 2 minutes, followed by 30 cycles of 98°C for 10s, 60°C for 30s, 72°C for 45s, and a final step of 72°C for 2 minutes. PCR products were run on a 1% gel at 100v for 60 minutes to determine the presence of targets.

##### S6. Library Preparation - GLST AmpSeq

##### First-Round PCR

An Illumina sequencing library was generated using a three-round PCR approach for samples obtained from 9 sites in Norway (n = 30) and 4 sites in Scotland (n = 28). Initially, samples were amplified in 55 individual reactions to amplify unique genomic targets per sample. Reaction conditions consisted of an initial incubation step of 98°C for 2 minutes, followed by 30 cycles of 98°C for 10s, 60°C for 30s, 72°C for 45s, and a final step of 72°C for 2 minutes at 10µl volumes (Q5 = 5µl, $dH_2O$ = 2.84µl, BSA = 0.16µl, forward primer = 0.5µl, reverse primer 0.5µl, and sample = 1µl). Single-target reactions from each sample were pooled into a 1.5ml Eppendorf with a working volume of ~820µl.

To remove first-round primer dimers, samples were run on a 1% agarose gel at 80V for 90 minutes. Each well contained 40µl of first-round PCR product and 16µl of loading dye and a 100bp ladder (New England Biolabs, Hitchin, Herts, UK) with a space between each sample. Bands were visualised under a transilluminator and cut out from the gel at the correct size (~300bp) according to the 100bp ladder. Gel slices were placed in a 1.5ml Eppendorf, 50µl of Nuclease-free $H_2O$ added, and left to incubate overnight at room temperature to pull the DNA out of the gel. The excised gel slice is removed, and the supernatant (Nuclease-free $H_2O$) is retained.

##### Second-Round PCR

Internal barcodes were added in a second-round PCR reaction with forward primer tag 5' – ACA CTC TTT CCC TAC ACG ACG CTC TTC CGA TCT AGG AGT CCT GAT GTT ATC CCT TGC ACC A – 3' (n = 61) and reverse primer tag 5' – GTG ACT GGA GTT CAG ACG TGT GCT CTT CCG ATC TAG GAG TCC CGT TGG AAG ATC GCG TCT AG – 3' (n = 62). Each 10µl barcoding reaction contained 5µl Q5 High-Fidelity Master Mix, 0.5µl of each forward and reverse barcode primer, and 4.5µl cleaned 1$^{st}$ round PCR product. The second-round PCR reaction conditions are 95˚C for 5 minutes, followed by 8 cycles of 95˚C for 30s, 60˚C for 30s, 72˚C for 1 minute, and a final extension step of 72˚C for 10 minutes. Products were run on a 1% agarose gel at 80V for 90 minutes to visualise a step-up in fragment size to confirm the addition of internal barcodes. Products were then gel extracted and purified using the same protocol as above to remove 2$^{nd}$ round and carry over 1$^{st}$ round primer dimers.

##### Third-Round PCR

A second barcoding reaction (third-round PCR) was conducted to add Illumina sequencing primer binding sites and identifier barcodes to each of the cleaned 2$^{nd}$ round PCR products. Each 20µl second barcoding reaction contained 10µl Q5 High-Fidelity Master Mix, 2µl of each forward and reverse barcode primer, 6µl $dH_2O$ and 2µl cleaned second-round PCR product. Reaction conditions were the same as the 2$^{nd}$ round of PCR. All samples (N = 58) were run on a 1% agarose gel at 80V for 90 minutes and graded by approximate concentration based on band intensity. The library was pooled by approximate volume based on grading to achieve approximate equimolar concentrations. Pooled library was run on a 1% agarose gel, band excised, and extracted using PureLink Quick Gel Extraction Kit (K210012, Invitrogen, Carlsbad, CA, USA) following the manufacturer's instructions. To remove any remaining impurities from the final extract library, an ethanol precipitation was carried out by adding 0.1 volume of 3M sodium acetate and 3 volumes of ice-cold 100% ethanol, followed by thorough mixing. The library was precipitated at -20˚C overnight and then centrifuged at 13,000rpm for 30 min at 4˚C the following day. The resultant pellet was washed twice with 500µl ice-cold 75% ethanol (10 min centrifugation at 4˚C per wash), briefly centrifuged for 10s at full speed to remove residual ethanol, and air-dried. The library was resuspended in 20µl nuclease-free water and quantified through the Nanodrop spectrophotometer (NanoDrop Ultra[C], Thermo Fisher Scientific, Waltham, MA, USA). The cleaned pooled library was subsequently sent to Genewiz® (Azenta Life Sciences) for high-throughput sequencing on a NovaSeq X Plus machine, 2 x 150bp paired-end reads.

#### S7. Demultiplexing and Mapping reads

Sequences from each sample were first processed to remove orphan reads using fastq_pair and then mapped to the draft reference genome using BWA **(Li and Durbin, 2009)**. The resultant sam files were then converted to BAM files, sorted and indexed using Samtools **(Li *et al.*, 2009)**. BAM files were then converted to .bed files in Bedtools **(Quinlan and Hall, 2010)** before being sorted and reads mapped to each genomic contig counted using cut and uniq commands in Linux. The resultant data were then plotted for each sample in R to assess mapping efficiency across each marker using **ggplot2**, **dplyr**, and **tidyr (Posit Software, 2024; R Core Team, 2024; Wickham, Vaughan and Girlich, 2024; Wickham *et al.*, 2023; Wickham, 2016)**.

#### S8. Phylogenetic Analysis

##### Variant Calling and Quality Control

Raw reads from the 58 Scottish and Norwegian isolates were mapped to the *Neoparamoeba* and *Perkinsela* draft reference assemblies using BWA-MEM **(Li, 2013)**. Joint variant calling was executed across all samples using **bcftools mpileup** and **call** under a multiallelic model **(Danecek *et al.*, 2021)**. While the inherent challenges of environmental sequencing resulted in a low median genome-wide coverage (~2–4x), we applied a highly stringent, multi-tiered filtering pipeline to isolate a panel of ultra-high-confidence markers. Individual genotype calls supported by fewer than three reads were masked as missing data (./.). Subsequently, variant sites were aggressively filtered to retain only high-quality single-nucleotide polymorphisms (QUAL=30) genotyped in at least 70% of the population (missingness < 0.30). This rigorous approach effectively stripped away low-coverage background noise, yielding a final targeted panel of 50 strictly biallelic host SNPs and 44 symbiont SNPs exhibiting deep sequencing coverage (mean site depth > 100x per isolate).

##### Population Genomic and Comparative Diversity Analyses

Filtered VCFs were imported into R Studio (v4.3.0) and converted to *genlight* and *genind* objects using **vcfR (Knaus & Grunwald, 2017)**. Pairwise fixation indices *($F_{ST}$)* were calculated using **StAMPP** (1000 bootstraps) **(Pembleton, Cogan & Foster, 2013)**. To visualise population structure and individual membership probabilities, Discriminant Analysis of Principal Components (**DAPC**) was performed using **adegenet (Jombart & Ahmed, 2011; Jombart, Devillard & Balloux, 2010)**, retaining A-score optimised principal components (1 PC for host, 5 PCs for symbiont) to prevent overfitting. Individual membership probabilities to geographic clusters were extracted and visualised as admixture proportions. Geographic **isolation-by-distance (IBD)** was assessed using Mantel tests (999 permutations) to correlate Euclidean geographic distances with Slatkin's linearised genetic distances $[F_{ST}/(1-F_{ST})]$ **(Legendre & Legendre, 2012)**.

#### Clonality and Linkage Disequilibrium

To account for potential sequencing artifacts and paralogous mapping noise, datasets were collapsed into Multi-Locus Genotypes (MLGs) using a 5% bitwise distance threshold in **poppr** **(Kamvar, Tabima & Grunwald, 2014)**. The reproductive mode of each partner was evaluated using the Index of Association $(I_A, rBar_d)$, with significance determined via 999 permutations against a null model of panmixia. To quantify the magnitude of genomic linkage, genome-wide pairwise linkage disequilibrium (LD) was calculated as $r^2$ for all biallelic SNPs using SNPRelate **(Zheng *et al.*, 2012)**. LD distributions were visualised using kernel density estimates, and the proportion of SNP pairs exceeding a biological threshold of $r^2 > 0.2$ was utilised as a proxy for clonal persistence. Genetic relationships between MLGs were further visualised using Minimum Spanning Networks (MSNs) based on bitwise genetic distances.

#### Cophylogenetic Congruence and Transmission Dynamics

Host-symbiont co-evolutionary signals were evaluated using several statistical approaches. Global genetic independence was tested via a Mantel test correlating host and symbiont distance matrices. We also employed Procrustes Approach to Cophylogeny (PACo) to assess topological congruence using the **vegan** and **paco** packages **(Oksanen *et al.*, 2025; Hutchinson *et al.*, 2017)**, and ParaFit analyses **(Legendre *et al.*, 2002)**. PACo was performed using 999 permutations to calculate global goodness-of-fit $(m^2)$, and jackknifed squared residuals were extracted to identify specific host-symbiont pairs exhibiting high phylogenetic incongruence. Fine-scale transmission dynamics were further visualised using a bipartite association network of host and symbiont MLGs constructed with **ggraph** and **igraph**, where edges were weighted by pairing frequency and coloured by geographic origin.

#### References


Amos, B., Aurrecoechea, C., Barba, M., Barreto, A., Basenko, E.Y., Bażant, W., Belnap, R., Blevins, A.S., Böhme, U., Brestelli, J., Brunk, B.P., Caddick, M., Callan, D., Campbell, L., Christensen, M.B., Christophides, G.K., Crouch, K., Davis, K., DeBarry, J., Doherty, R., Yikun, D., Dunn, M., Falke, D., Fisher, S., Flicek, P., Fox, B., Gajria, B., Giraldo-Calderón, G.I., Harb, O.S., Harper, E., Hertz-Fowler, C., Hickman, M.J., Howington, C., Hu, S., Humphrey, J., Iodice, J., Jones, A., Judkins, J., Kelly, S.A., Kissinger, J.C., Kun Kwon, D., Lamoureux, K., Lawson, D., Li, W., Lies, K., Lodha, D., Long, J., MacCallum, R.M., Maslen, G., McDowell, M.A., Nabrzyski, J., Roos, D.S., Rund, S.S.C., Schulman, S.W., Shanmugasundram, A., Sitnik, V., Spruill, D., Stoeckert, C.J., Tomko, S.S., Wang, H., Warrenfeltz, S., Wieck, R., Wilkinson, P.A., Xu, L. and Zheng, J. (2022) 'VEuPathDB: the eukaryotic pathogen, vector and host bioinformatics resource center', *Nucleic Acids Research*, 50(D1), pp. D898–D911. Available at: https://doi.org/10.1093/nar/gkab929

Andrews, S. (2010) *FastQC: A Quality Control Tool for High Throughput Sequence Data*. Babraham Bioinformatics. Available at: http://www.bioinformatics.babraham.ac.uk/projects/fastqc/

Bushnell, B. (2014). BBMap: A Fast, Accurate, Splice-Aware Aligner. Lawrence Berkeley National Laboratory.

Danecek, P. et al. (2021) ‘Twelve years of SAMtools and BCFtools’, *GigaScience*, 10(2), p. giab008. Available at: https://doi.org/10.1093/gigascience/giab008

De Coster, W., D’Hert, S., Schultz, D.T., Cruts, M. and Van Broeckhoven, C. (2018) ‘NanoPack: visualizing and processing long-read sequencing data’, *Bioinformatics*, 34(15), pp. 2666–2669. Available at: https://doi.org/10.1093/bioinformatics/bty149

Galili, T. (2015) ‘dendextend: an R package for visualizing, adjusting and comparing trees of hierarchical clustering’, *Bioinformatics*, 31(22), pp. 3718–3720. Available at: https://doi.org/10.1093/bioinformatics/btv428

Hutchinson, M.C. et al. (2017) ‘Paco: implementing Procrustean Approach to Cophylogeny in R’, *Methods in Ecology and Evolution*, 8(8), pp. 932–940. Available at: https://doi.org/10.1111/2041-210X.12736

Jombart, T., Devillard, S. and Balloux, F. (2010) ‘Discriminant analysis of principal components: a new method for the analysis of genetically structured populations’, *BMC Genetics*, 11(1), p. 94. Available at: https://doi.org/10.1186/1471-2156-11-94

Jombart, T. and Ahmed, I. (2011) ‘adegenet 1.3-1: new tools for the analysis of genome-wide SNP data’, *Bioinformatics*, 27(21), pp. 3070–3071. Available at: https://doi.org/10.1093/bioinformatics/btr521

Kamvar, Z.N., Tabima, J.F. and Grünwald, N.J. (2014) ‘Poppr: an R package for genetic analysis of populations with mixed reproduction’, *PeerJ*, 2, p. e281. Available at: https://doi.org/10.7717/peerj.281

Knaus, B.J. and Grünwald, N.J. (2017) ‘vcfr: a package to manipulate and visualize variant call format data in R’, *Molecular Ecology Resources*, 17(1), pp. 44–53. Available at: https://doi.org/10.1111/1755-0998.12549

Kolmogorov, M., Yuan, J., Lin, Y. and Pevzner, P.A. (2019) ‘Assembly of long, error-prone reads using repeat graphs’, *Nature Biotechnology*, 37(5), pp. 540–546. Available at: https://doi.org/10.1038/s41587-019-0072-8

Kriventseva, E.V., Kuznetsov, D., Tegenfeldt, F., Manni, M., Dias, R., Simão, F.A. and Zdobnov, E.M. (2019) ‘OrthoDB v10: sampling the diversity of animal, plant, fungal, protist, bacterial and viral genomes for evolutionary and functional annotations of orthologs’, *Nucleic Acids Research*, 47(D1), pp. D807–D811. Available at: https://doi.org/10.1093/nar/gky1053

Legendre, P. and Legendre, L. (2012) *Numerical Ecology*. 3rd edn. Amsterdam: Elsevier.

Legendre, P., Desdevises, Y. and Bazin, E. (2002) ‘A statistical test for host–parasite coevolution’, *Systematic Biology*, 51(2), pp. 217–234. Available at: https://doi.org/10.1080/10635150252899734

Li, H. (2013) ‘Aligning sequence reads, clone sequences and assembly contigs with BWA-MEM’, *arXiv*. Available at: https://arxiv.org/abs/1303.3997

Li, H. (2018) ‘Minimap2: pairwise alignment for nucleotide sequences’, *Bioinformatics*, 34(18), pp. 3094–3100. Available at: https://doi.org/10.1093/bioinformatics/bty191

Li, H. and Durbin, R. (2009) ‘Fast and accurate short read alignment with Burrows–Wheeler transform’, *Bioinformatics*, 25(14), pp. 1754–1760. Available at: https://doi.org/10.1093/bioinformatics/btp324

Li, H., Handsaker, B., Wysoker, A., Fennell, T., Ruan, J., Homer, N., Marth, G., Abecasis, G. and Durbin, R. (2009) ‘The Sequence Alignment/Map format and SAMtools’, *Bioinformatics*, 25(16), pp. 2078–2084. Available at: https://doi.org/10.1093/bioinformatics/btp352

Li, D., Liu, C.M., Luo, R., Sadakane, K. and Lam, T.W. (2015) ‘MEGAHIT: an ultra-fast single-node solution for large and complex metagenomics assembly via succinct de Bruijn graph’, *Bioinformatics*, 31(10), pp. 1674–1676. Available at: https://doi.org/10.1093/bioinformatics/btv033

Manni, M., Berkeley, M.R., Seppey, M., Simão, F.A. and Zdobnov, E.M. (2021) ‘BUSCO update: novel and streamlined workflows along with broader and deeper phylogenetic coverage for scoring of eukaryotic, prokaryotic, and viral genomes’, *Molecular Biology and Evolution*, 38(10), pp. 4647–4654. Available at: https://doi.org/10.1093/molbev/msab199

Oksanen, J. et al. (2024) *vegan: Community Ecology Package*. R package version 2.7-3. Available at: https://CRAN.R-project.org/package=vegan

Pebesma, E. (2018) ‘Simple Features for R: standardized support for spatial vector data’, *The R Journal*, 10(1), pp. 439–446. Available at: https://doi.org/10.32614/RJ-2018-009

Pembleton, L.W., Cogan, N.O.I. and Forster, J.W. (2013) ‘StAMPP: an R package for calculation of genetic differentiation and structure of mixed-ploidy level populations’, *Molecular Ecology Resources*, 13(5), pp. 946–952. Available at: https://doi.org/10.1111/1755-0998.12129

Posit Software, PBC. (2024) *RStudio: Integrated Development Environment for R*. Version 2024.09.0 Build 375 “Cranberry Hibiscus”. Available at: https://posit.co/

Quinlan, A.R. and Hall, I.M. (2010) ‘BEDTools: a flexible suite of utilities for comparing genomic features’, *Bioinformatics*, 26(6), pp. 841–842. Available at: https://doi.org/10.1093/bioinformatics/btq033

R Core Team (2024) *R: A Language and Environment for Statistical Computing*. Vienna: R Foundation for Statistical Computing. Available at: https://www.R-project.org/

Saary, P., Mitchell, A.L. and Finn, R.D. (2020) ‘Estimating the quality of eukaryotic genomes recovered from metagenomic analysis with EukCC’, *Genome Biology*, 21(1), p. 244. Available at: https://doi.org/10.1186/s13059-020-02155-4

Schmieder, R. and Edwards, R. (2011) ‘Fast identification and removal of sequence contamination from genomic and metagenomic datasets’, *PLoS ONE*, 6(3), p. e17288. Available at: https://doi.org/10.1371/journal.pone.0017288

Schwabl, P., Maiguashca Sánchez, J., Costales, J.A., Ocaña-Mayorga, S., Segovia, M., Carrasco, H.J., Hernández, C., Ramírez, J.D., Lewis, M.D., Grijalva, M.J. and Llewellyn, M.S. (2020) ‘Culture-free genome-wide locus sequence typing (GLST) provides new perspectives on *Trypanosoma cruzi* dispersal and infection complexity’, PLOS Genetics, 16(12), p. e1009170. Available at: https://doi.org/10.1371/journal.pgen.1009170

Seemann, T. (2013) *any2fasta: Convert Various Sequence Formats to FASTA*. GitHub repository. Available at: https://github.com/tseemann/any2fasta

Slowikowski, K. (2023) *ggrepel: Automatically Position Non-Overlapping Text Labels with "ggplot2"*. R package version 0.9.4. Available at: https://CRAN.R-project.org/package=ggrepel

South, A., Michael, S. and Massicotte, P. (2023) *rnaturalearthdata: World Vector Map Data from Natural Earth Used in "rnaturalearth"*. R package version 0.1.0. Available at: https://CRAN.R-project.org/package=rnaturalearthdata

Stanke, M. and Morgenstern, B. (2005) ‘AUGUSTUS: a web server for gene prediction in eukaryotes’, *Nucleic Acids Research*, 33(suppl_2), pp. W465–W467. Available at: https://doi.org/10.1093/nar/gki458

Stanke, M., Diekhans, M., Baertsch, R. and Haussler, D. (2008) ‘Using native and syntenically mapped cDNA alignments to improve de novo gene finding’, *Bioinformatics*, 24(5), pp. 637–644. Available at: https://doi.org/10.1093/bioinformatics/btn013

Wickham, H. (2016) *ggplot2: Elegant Graphics for Data Analysis*. New York: Springer-Verlag.

Wickham, H., François, R., Henry, L., Müller, K. and Vaughan, D. (2023) *dplyr: A Grammar of Data Manipulation*. R package version 1.1.4. Available at: https://dplyr.tidyverse.org

Wickham, H., Vaughan, D. and Girlich, M. (2024) *tidyr: Tidy Messy Data*. R package version 1.3.1. Available at: https://tidyr.tidyverse.org

Young, N.D., Dyková, I., Snekvik, K., Nowak, B.F. and Morrison, R.N. (2008) ‘Neoparamoeba perurans is a cosmopolitan aetiological agent of amoebic gill disease’, *Diseases of Aquatic Organisms*, 78(3), pp. 217–223. Available at: https://doi.org/10.3354/dao01869

Zimin, A.V., Marçais, G., Puiu, D., Roberts, M., Salzberg, S.L. and Yorke, J.A. (2013) ‘The MaSuRCA genome assembler’, *Bioinformatics*, 29(21), pp. 2669–2677. Available at: https://doi.org/10.1093/bioinformatics/btt476
