## Supplemental Figure 1, Table 1, and Figure 2. Figure one is not referenced in the text - to illustrate reads per loci per site. for "Asymmetric population structure in *Neoparamoeba perurans* and its kinetoplastid symbiont"

### Supplementary Data 2 (SD2) – Results

#### S1. Target recovery and sequencing coverage across sampling sites

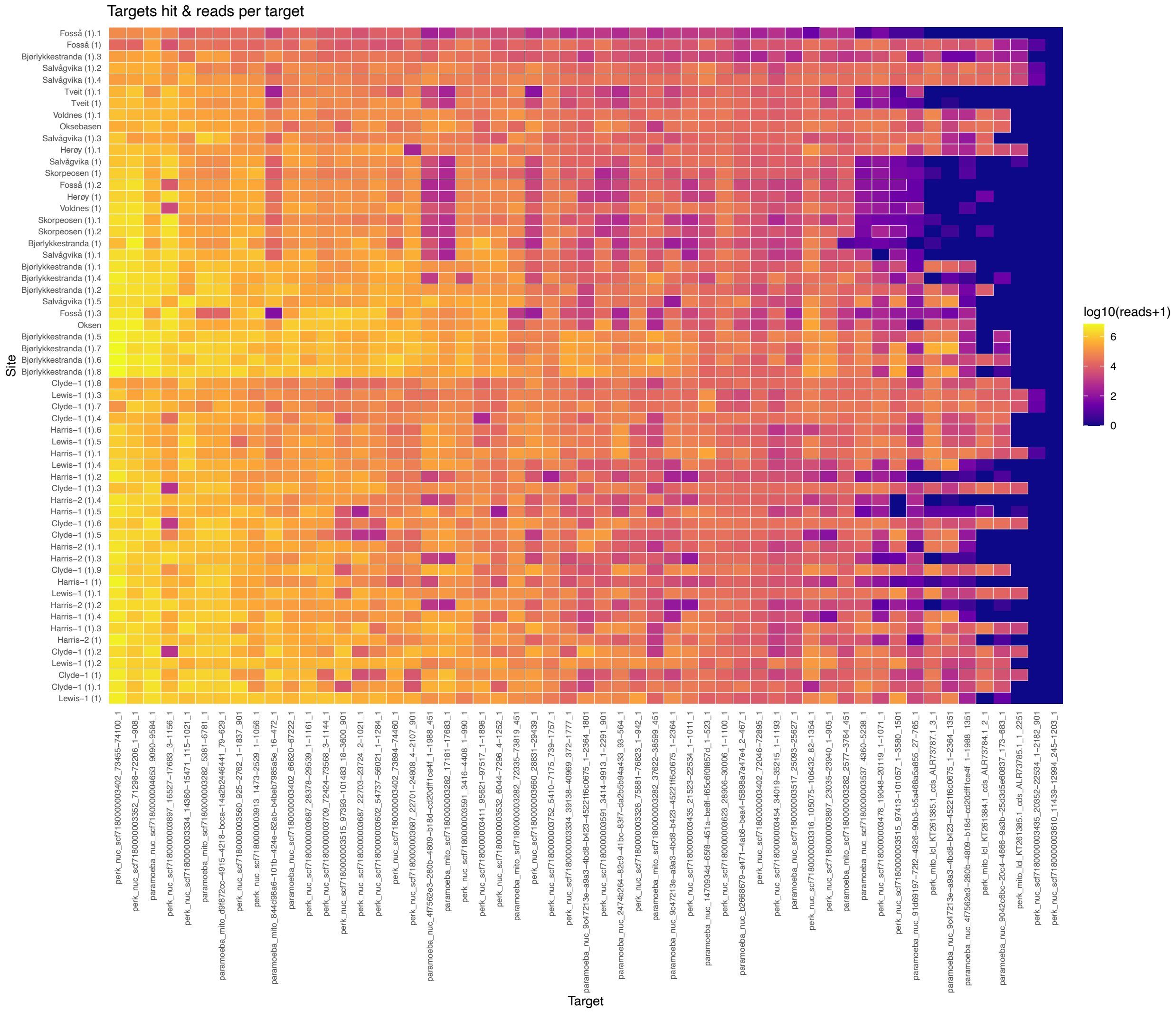


**Figure S1: Heatmap of target locus recovery and sequencing coverage across sampling sites.** Rows represent sampling site locations and columns represent target loci. Cell colour indicates the number of sequencing reads mapped to each target locus, with warmer colours representing higher read coverage and cooler colours indicating lower coverage or target dropout. Variation in target recovery was observed among loci and sampling sites, highlighting differences in sequencing performance and locus representation across the dataset.

#### S2. Comparative genomic metrics for *Neoparamoeba* and *Perkinsela*

**Table S1: Comparative genomic metrics for *Neoparamoeba* and its endosymbiont *Perkinsela*.** *Denotes statistical significance at $p < 0.05$. Note: G/N represents the ratio of unique genotypes to total samples, a common measure of clonal richness.*

| Metric Category | Genetic Parameter | *Neoparamoeba* (Host) | *Perkinsela* (Endosymbiont) |
|---|---|---|---|
| Sample Summary | SNP Count (Filtered) | 50 | 44 |
| Population Structure | Global Fixation Index ($F_{ST}$) | 0.008 *(p = 0.123)* | 0.035* *(p < 0.001)* |
| | Isolation By Distance (IBD) Correlation (Mantel *r)* | *0.09* (p = 0.001)* | *0.12* (p = 0.001)* |
| Reproductive Mode | Index of Association ($rBar_d$) | 0.0421* *(p = 0.001)* | 0.254* *(p = 0.001)* |
| | Pairwise Linkage Disequilibrium (LD) ($r^2 > 0.2$) | *4.84%* | *17.49%* |
| Clonal Diversity | Unique Multi-Locus Genotypes (MLGs) | 47 | 21 |
| | Genotypic Diversity *(G/N)* | 0.810 | 0.362 |
| Cophylogeny | Mantel Host-Symbiont *(r)* | *0.130* (p = 0.049)* | |
| | PACo Global Fit ($m^2$) | *14.3* (p < 0.001)* | |
| | ParaFit Global Stat | *5.23* (p = 0.039)* | |

#### S3. Clonality and cophylogenetic analyses

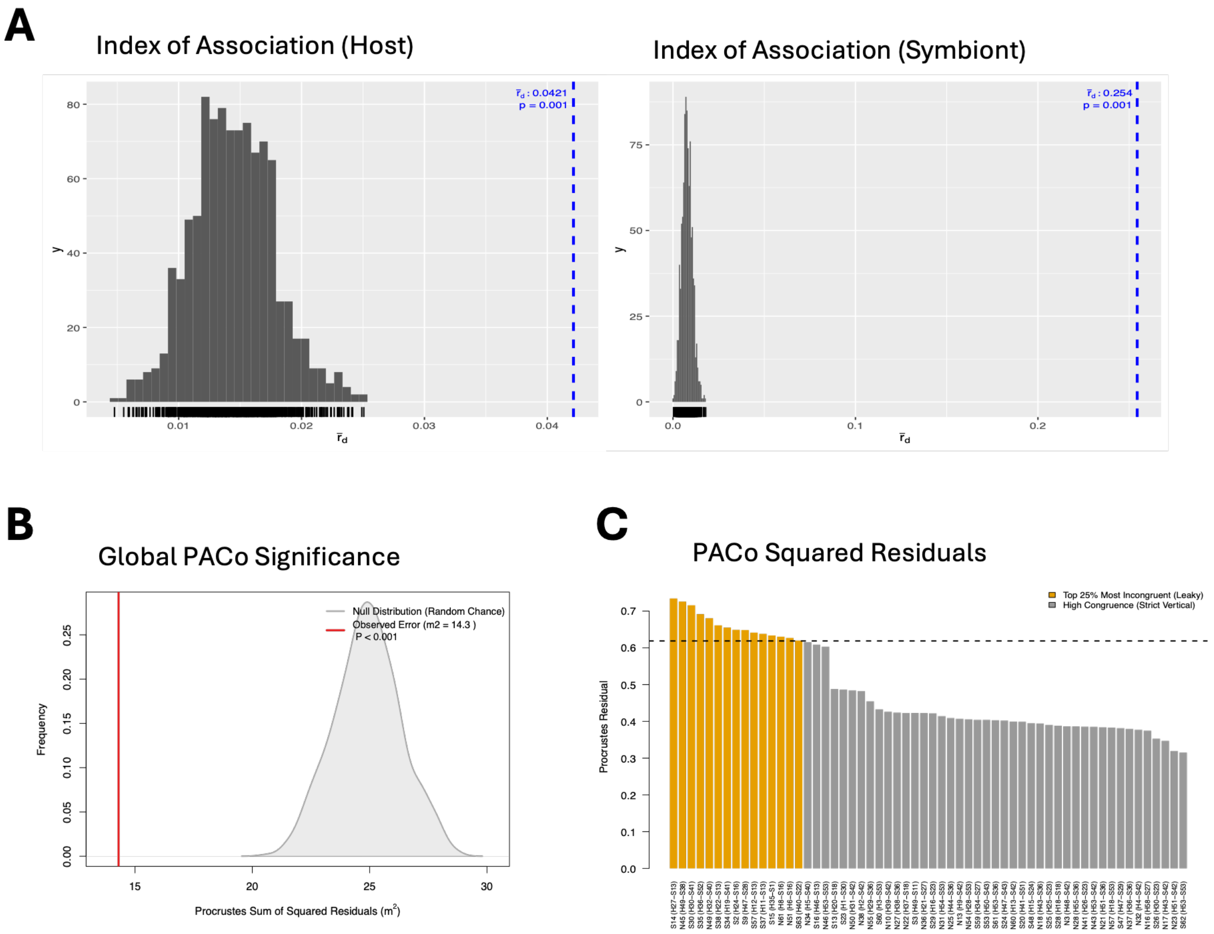


**Figure S2: Analysis of clonality and host–symbiont cophylogenetic congruence. (A) Distribution of the standardised Index of Association ($\bar{r}_d$) following 999 permutations for host and symbiont datasets.** Observed values (blue dashed lines) fall outside the null distributions generated under random association *($P_{host}$ = 0.001, $P_{symbiont}$ = 0.001)*, supporting significantly clonal reproductive modes under outbreak conditions. **(B) Global PACo significance test.** The vertical red line indicates the observed Procrustes sum of squared residuals ($m^2 = 14.3$) plotted against the null distribution generated from 999 permutations. An observed error significantly lower than expected under the null model indicates significant global congruence between host and symbiont topologies ($P < 0.001$), supporting a primary mode of vertical transmission. **(C) Distribution of PACo Procrustes squared residuals for individual host–symbiont associations.** Bars represent individual isolate pairings ranked by residual magnitude; orange bars indicate the top 25% most incongruent associations (residuals above the 75th percentile), highlighting candidate cases of horizontal transmission or "leaky" inheritance. High-incongruence outliers include isolates such as S9 and N51.
